# Adiponectin Signaling Regulates Urinary Bladder Function by Blunting Smooth Muscle Purinergic Contractility

**DOI:** 10.1101/2024.10.25.620328

**Authors:** Zhaobo Luo, Ali Wu, Simon Robson, Seth Alper, Weiqun Yu

## Abstract

Lower urinary tract symptoms (LUTS) affect ∼ 50% of the population aged >40 years and are strongly associated with obesity and metabolic syndrome. Adipose tissue plays a key role in obesity/metabolic syndrome by releasing adipokines that regulate systemic energy/lipid metabolism, insulin resistance, and inflammation. Adiponectin (ADPN), the most abundant adipokine, modulates energy/metabolism homeostasis through its insulin-sensitizing and anti-inflammatory effects. Human plasma ADPN levels are inversely associated with obesity and diabetes. The role of adipokines such as ADPN in the LUTS associated with obesity/metabolic syndrome remains unknown. We have tested such a possible role in a global ADPN knockout mouse model (*Adpn^−/−^*). *Adpn^−/−^* mice exhibited increased voiding frequency, small voids, and reduced bladder smooth muscle (BSM) contractility with absence of purinergic contraction. Molecular examination indicated significantly altered metabolic and purinergic pathways. The ADPN receptor agonist AdipoRon was found to abolish acute BSM contraction. Intriguingly, both AMPK activators and inhibitors also abolished BSM purinergic contraction. These data indicate the important contribution of a novel ADPN signaling pathway to the regulation of BSM contractility. Dysregulation of this ADPN signaling pathway might be an important mechanism leading to LUTS associated with obesity/metabolic syndrome.

**ARTICLE HIGHLIGHTS:** Lower urinary tract symptom (LUTS) is strongly associated with obesity and metabolic syndrome, however, the underlying molecular mechanisms are missing. Dysregulation of adipokine signaling could be the link for this association.

Whether adiponectin, the most abundant adipokine, plays a role in regulating bladder function and dysfunction.

Mice null for adiponectin exhibited increased voiding frequency, small voids, and reduced bladder smooth muscle contractility, with corresponding metabolic and purinergic pathway changes.

Dysregulation of adiponectin signaling might be an important mechanism leading to LUTS associated with obesity/metabolic syndrome.

## Introduction

Lower urinary tract symptoms (LUTS) such as urinary incontinence (UI), urgency, overactive bladder (OAB) and underactive bladder (UAB) affect ∼ 50% of the population aged 40 and over (1; 2). LUTS is strongly associated with obesity and metabolic syndrome (3–6). Obesity affects over one-third of the US population and is clinically recognized as a predisposing risk factor to both UI and OAB. Obesity is also a major risk factor for type II diabetes, hypertension, dyslipidemia, and atherosclerosis, together described as metabolic syndrome. Visceral obesity is considered a hallmark of metabolic syndrome, which promotes cellular dysfunction and accelerates aging (7–9). Diabetes mellitus afflicts 10.5% of the US population (National Health and Nutrition Examination Survey, 2020), of whom at least 40% (and in some studies up to 100%) experience diabetic bladder dysfunction (DBD) (10; 11). A direct link between obesity/metabolic syndrome and LUTS is suggested by the beneficial effects of bariatric surgery, caloric restriction, and physical exercise on both UI and OAB (3; 4; 6; 12; 13).

The underlying molecular mechanisms by which obesity and metabolic syndrome predispose to LUTS are poorly understood. Hyperglycemic or ischemic oxidative stress or chronic low-grade inflammation due to metabolic stress might contribute to LUTS pathogenesis (14). However, recent advances in our understanding of energy balance and metabolic syndrome provide important hints on potential molecular links between LUTS and obesity/metabolic syndrome. Adipose tissue (fat), in addition to functioning as an energy storage site, is also an important endocrine organ releasing physiologically active, circulating adipokines, including leptin, tumor necrosis factor α (TNF-α), adipsin, plasminogen activator inhibitor-1, interleukin 6, resistin, monocyte chemotactic protein-1, adiponectin (ADPN) and others. These adipokines are important regulators of energy and lipid metabolism, insulin resistance, inflammation, and many other cellular functions (7-9; 15). However, the relationship between adipokines and LUTS associated with obesity/metabolic syndrome remains unclear.

We have recently reported that mice with smooth muscle-specific deletion of the insulin receptor (*SMIR^−/−^*) had abnormal bladder phenotypes, including urinary frequency and small voids, and decreased contractility of bladder smooth muscle (BSM). The study suggested that dysregulated insulin signaling or insulin resistance in local bladder tissue might be important to the development of DBD (16). Interestingly, we observed downregulation of ADPN expression in bladders of *SMIR^−/−^*mice (16). As ADPN is predominantly released by adipose tissue, we investigated bladder tissue expression of ADPN. Our immunofluorescence localization data indicated the expression of ADPN in BSM cells (16). These discoveries suggest a novel ADPN signaling mechanism contributing to the regulation of bladder function.

ADPN is the most abundant adipokine secreted mainly by adipose tissue (17; 18). ADPN levels decrease with progression of obesity/metabolic syndrome and with age, whereas elevated levels are generally protective (19–22). Elevated ADPN levels are believed to exert insulin-sensitizing, anti-diabetic, anti-inflammatory, and anti-atherogenic effects. Global ADPN knockout mice fed a high-fat diet develop exacerbated insulin resistance and metabolic syndrome, with shortened lifespan (23–25). In contrast, transgenic mice with elevated circulating ADPN levels exhibit dramatically improved systemic insulin sensitivity, reduced age-related tissue inflammation, and prolonged lifespan (24). ADPN gene mutations and polymorphisms in humans are closely associated with low circulating ADPN levels, insulin resistance, and type II diabetes (26; 27). These studies suggest ADPN as a key adipokine modulating progression of metabolic syndrome. Whether and how adipokine signaling regulates bladder function has remained unclear. However, in a mouse model with a 56% increase in plasma ADPN, BSM cells exhibited altered Ca^2+^ sensitivity, supporting a functional role of ADPN signaling in BSM (28).

The association between LUTS and obesity/metabolic syndrome, and the strong correlation between adipokine dysregulation and obesity/metabolic syndrome in both patients and animal models prompted our hypothesis that dysregulated adipokine signaling is a major contributor to the pathogenesis of LUTS associated with obesity/metabolic syndrome. We have tested our hypothesis in this study using a global ADPN knockout mouse model.

## Methods and materials

### Reagents

Unless otherwise specified, all chemicals were obtained from MilliporeSigma and were of reagent grade or better. P2X1 receptor agonist α,β-methyleneadenosine 5’-triphosphate trisodium (α,β-meATP, Catalog #: 3209), muscarinic receptor agonist carbachol (Catalog #: 2810), and ADPN receptor agonist AdipoRon (Catalog #: 5096) were purchased from R&D Systems. AMP-activated protein kinase (AMPK) inhibitor Bay3827 (Catalog #: HY-112083), AMPK activator A769662 (Catalog #: HY-50662), and AMPK activator EX229 (Catalog #: HY-112769) were purchased from MedChemExpress.

### Animals

All mice used in this study were of C57BL/6J background and matched for sex and age (12–16 weeks). Both male and female mice were used unless otherwise specified. Mice were housed in standard polycarbonate cages with free access to normal food and water. Wild-type C57BL/6J mice and global ADPN knockout mice (*Adpn^−/−^*, strain #: 008195) were purchased from Jackson Laboratory (Bar Harbor, ME). All animal studies were performed in adherence to U.S. National Institutes of Health guidelines for animal care and use, and with approval of the Beth Israel Deaconess Medical Center Institutional Animal Care and Use Committee.

### Void spot assay (VSA)

VSA was performed during the daytime with light from ∼ 9:00 to 13:00 in both male and female mice. Individual mice were gently placed in a standard mouse cage lined with Blicks Cosmos Blotting Paper (Catalog no. 10422-1005), without drinking water but with standard dry mouse chow available. Blotting paper was recovered and imaged as described (29). Overlapping voiding spots were visually examined and manually separated by outlining and copying, then pasting to a nearby empty space in Fiji software for analysis. A volume:area standard curve defined a 1-mm^2^ voiding spot as representing 0.283 μL of urine. Urine spots with an area ≥ 80 mm^2^ were considered primary voiding spots (PVS) based on a cutoff established from the voiding spot patterns of hundreds of mice (29; 30).

### Cystometrogram (CMG)

CMG performed in *wild-type* female mice with PBS infusion (25 μL/min) as previously described produces a consistent voiding pattern (31; 32). With urethane (1.4 g/kg body weight) and continuous-flow isoflurane (3% induction, 1.0% maintenance) anesthetization, a 1-cm midline abdominal incision was performed. PE50 tubing was then implanted through the dome of the bladder. The catheter was connected to a pressure transducer and a syringe pump was coupled to data-acquisition devices (WPI Transbridge and AD Instruments PowerLab 4/35) and a computerized recording system (AD Instruments LabChart software). Isoflurane was withdrawn immediately after surgery and CMG was performed under urethane anesthesia, which spares the voiding reflex. CMG was performed for > 1 hour of stable, regular voiding cycles. At least 5 cycles of filling and voiding traces were assessed to evaluate urodynamics. Bladder compliance was determined by 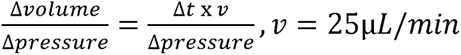.

### BSM Myography

BSM myography was performed on male mice, whose ∼30 mg bladders (larger than in females) facilitate dissection of the bladder epithelial layer away from BSM for a consistent per-bladder yield of 4 ∼7 mm by ∼2 mm muscle strips free of mucosa. Muscle strips were mounted in an SI-MB4 tissue bath system (World Precision Instruments, FL, USA). Force sensors were connected to a TBM 4M transbridge (World Precision Instruments). Signals were amplified by PowerLab (AD Instruments, CO, USA) and monitored with Chart software (AD Instruments). BSM strips gently stretched to optimize contractile force were then pre-equilibrated for at least 1 h. Contraction force was sampled at 2000/s using Chart software. Electrical field stimulation (EFS) was carried out using a Grass S48 field stimulator (Grass Technologies, RI, USA) using previously described standard protocols: voltage 50 V; duration: 0.05 ms; trains of stimuli: 3 s; frequencies: 1, 2, 5, 10, 20 and 50 Hz (31; 32).

### Immunofluorescence staining and imaging

Excised bladders fixed in 4% (w/v) paraformaldehyde were cryoprotected, frozen, sectioned, and incubated overnight at 4 °C with antibodies (1:100, Supplemental Table 1). The sections were then incubated with an Alexa Fluor 488–conjugated secondary antibody (diluted 1:100), and nuclei were stained with DAPI. Imaging was performed on an Olympus BX60 fluorescence microscope with a 40×/0.75 objective (32).

### Western blot

Whole bladder tissues from female mice were lysed with RIPA buffer (50 mmol/L Tris-HCl, 150 mmol/L NaCl, 1% NP-40, 0.5% sodium deoxycholate, and 0.1% SDS). Protein concentrations were determined by BCA protein assay (Thermo Scientific, Rockford, IL, USA). 25 µg protein per well was fractionated by SDS-PAGE and transferred to PVDF membrane. Membranes were incubated with primary antibodies (Supplemental Table 1) at 4°C for 12 hours. Immunoreactive protein bands were visualized with Amersham ECL reagent (Arlington Heights, IL, USA). The membranes were incubated with Restore Plus Western Blot Stripping buffer (Thermo Fisher Scientific, Rockford, IL, USA) for 5 min to remove primary and secondary antibodies for reblotting with new antibodies, and each membrane was stripped and re-probed for no more than three times. Scanned protein band intensity was quantitated using Fiji software (32). The protein band intensity of interest was normalized to glyceraldehyde 3-phosphate dehydrogenase (GAPDH) intensity in the same lane. The average protein expression level in wild-type mice was designated as 1, and the fold change of protein expression level in knockout mice was reported.

### Statistical analyses

All data are presented as box and whisker plots or as individual symbols and line plots. The centerline in box and whisker plots is the median, the box represents 75% of the data, and the whiskers indicate minimum to maximum. Data were analyzed by 1-way ANOVA for comparison among groups. Data were analyzed by Student’s *t*-test between two groups. If possible, a paired *t*-test was used. Bonferroni’s multiple comparison post-hoc tests were used where necessary, and P < 0.05 was considered significant.

### Data and Resource Availability

Further information, reagents and other supporting data in this study are available from the corresponding author upon request.

## Results

### *Adpn^−/−^* mice exhibit altered voiding phenotype and urodynamics

We performed VSA to examine whether deletion of ADPN can directly cause voiding dysfunction in *Adpn^−/−^* mice (Fig. 1). Whereas wild-type mice produce ∼3 PVS over 4 h (PVS; male: 3.2 PVS/4; female: 3.5 PVS/4 h), mean PVS number in *Adpn^−/−^* mice was significantly higher (male: 4.9 PVS/4 h; female: 5.0 PVS/4 h). In contrast, *Adpn^−/−^* mean PVS area was significantly smaller (male: 94.2 μL/void; female: 85.1 μL/void) than for wild-type mice (male: 151.1 μL/void; female: 108.5 μL/void) (Fig. 1). To confirm the altered voiding pattern detected in VSA and to explore potential underlying mechanisms, urodynamic studies were performed by CMG. CMG data consistently indicated that *Adpn^−/−^*mice exhibited increased voiding frequency with decreased bladder compliance (Fig. 2).

**Figure 1.**
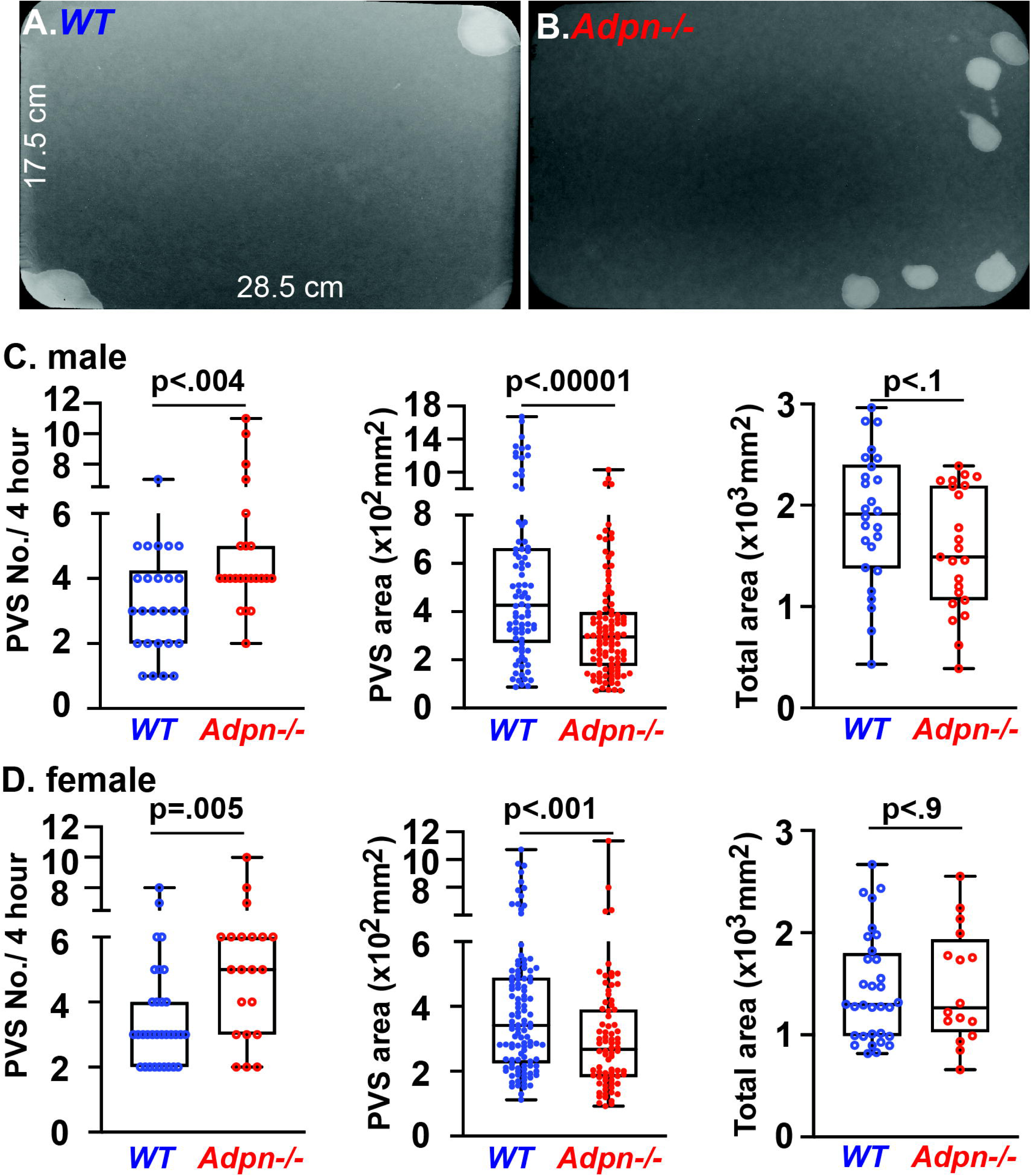
*Adpn^−/−^*mice exhibit increased voiding frequency and small voids. Representative VSA filter images are shown as *wild-type* (*A*: from n=26 male, n=32 female) and *Adpn^−/−^* (*B*: from n=23 male, n=21 female). *C* and *D*: Primary voiding spot counts (PVS) are defined as voiding spots > 80mm^2^ of males (*C*) and females (*D*). Data are plotted in Box (75% of the data) and whiskers format (minimum to maximum), with centerline as median value. Student *t*-test, with *P* values above bars.

**Figure 2.**
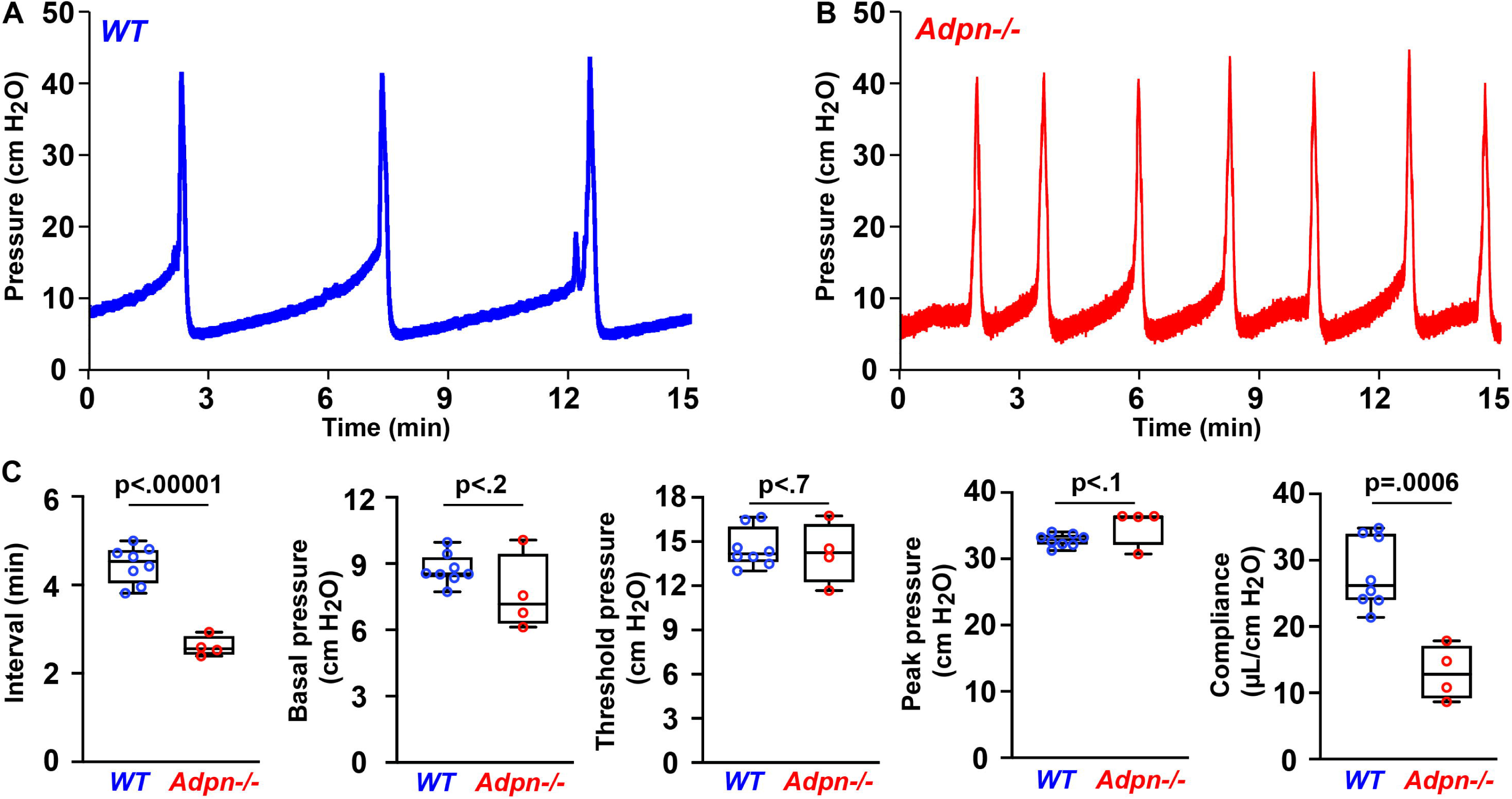
*Adpn^−/−^*mice exhibit voiding frequency with reduced compliance. Representative CMG traces of (*A*) *wild-type* (n=8) and (*B*) *Adpn^−/−^*mice (n=4) are quantitated in (*C*). Data are plotted in Box (75% of the data) and whiskers format (minimum to maximum), with centerline as median value. Student *t*-test, with *P* values above bars.

### *Adpn^−/−^* mice have diminished smooth muscle contraction force

BSM cells play a central role in bladder contraction and relaxation. As our previous study detected ADPN expression in BSM cells (16), we hypothesized that *Adpn* deletion could impair BSM contractility and alter bladder voiding phenotype in *Adpn^−/−^* mice (Fig. 1 & 2). Myography was performed on isolated BSM strips (Fig. 3*A* & *C*). BSM strip contractile force increased in response to increased EFS frequencies, mimicking the *in vivo* BSM contraction in response to neurotransmitter release. The contractile force of *Adpn^−/−^*mice BSM strips was significantly lower than for wild-type BSM strips (Fig. 3*A* & *C*). Wild-type bladder contraction is mainly mediated by parasympathetic ATP and Acetylcholine (ACh) co-release, which induces purinergic P2X1 and muscarinic CHRM3 receptor-mediated signaling cascades leading to BSM contraction (31). To measure possible changes of these signaling pathways in *Adpn^−/−^* mouse BSM, we tested the effect on BSM contraction of the CHRM3 receptor antagonist, atropine. As expected, atropine significantly inhibited wild-type mice BSM contraction force in response to EFS (Fig 3*B* & *D*), and the remaining contraction force (∼40%) is generally purinergic. To our surprise, atropine completely inhibited *Adpn^−/−^* mice BSM contraction in response to EFS (Fig.3*B* & *D*), suggesting a lack of purinergic contraction in *Adpn^−/−^* mouse bladder. Further studies using the P2X1 receptor agonist, α,β-meATP, confirmed minimal purinergic contractile force in *Adpn^−/−^* mouse bladder (Fig. 3*E*). To confirm the diminished contraction force in *Adpn−/−* mice bladder is intrinsic to BSM and not secondary to altered neurotransmitter release, we tested the BSM response to depolarization by high extracellular KCl. Results consistently indicated significantly smaller KCl-induced contraction force in BSM strips from *Adpn^−/−^* mice bladders (Fig. 4*F*).

**Figure 3.**
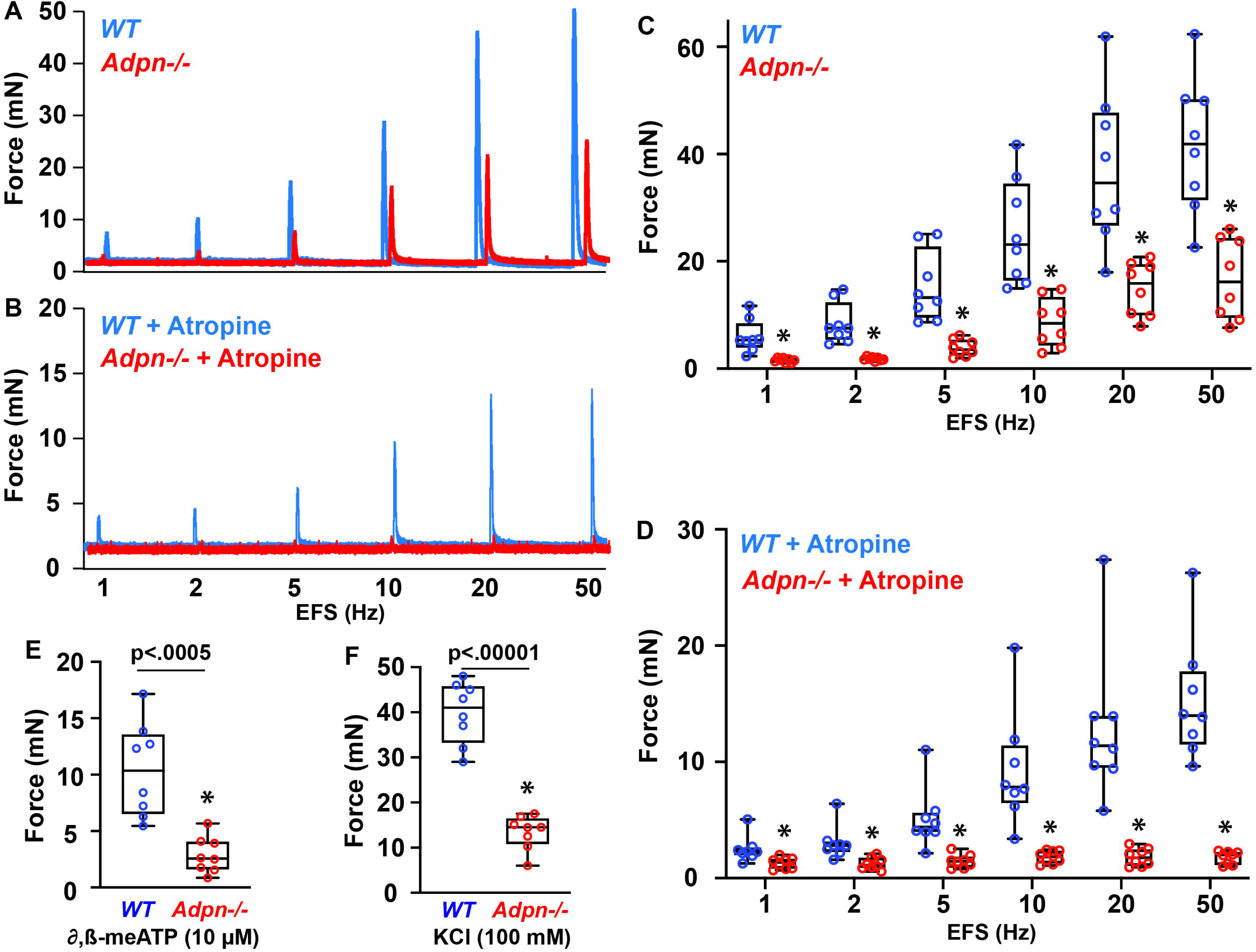
*Adpn^−/−^*BSM exhibits diminished contractile force, with absence of intrinsic purinergic contraction. Representative BSM contraction force traces in response to EFS are shown as *A* (Blue: *wild-type*, n=8; Red: *Adpn^−/−^,* n=8), and *B* in the presence of 0.5 µM atropine (Blue: *wild-type*, n=8; Red: *Adpn^−/−^,* n=8). *C* and *D*: Summarized data from multiple experiments in the format of *A* & *B*. *E* and *F*: BSM contractile force in response to α,β-meATP (*E*) and in response to KCl (*F*). Data are plotted in Box (75% of the data) and whiskers format (minimum to maximum), with centerline as median value. Student *t*-test, with *P* values above bars. *, p < 0.05.

**Figure 4.**
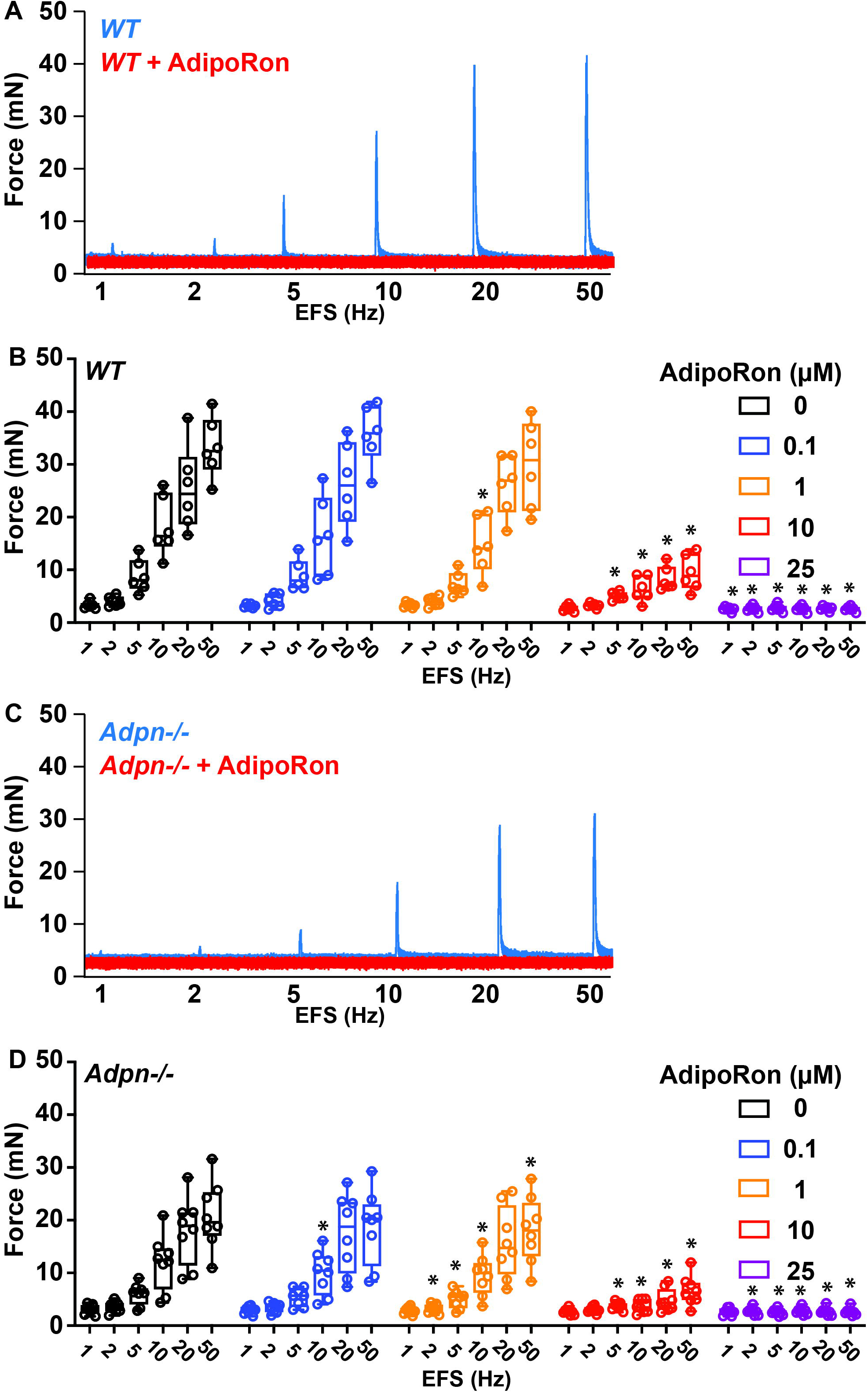
Adiponectin receptor agonist AdipoRon inhibits mouse BSM contractile force in a dose-dependent manner. *A* and *C*: Representative traces of male *wild-type* (n=6) and *Adpn−/−* (n=8) BSM contraction in response to EFS before (Blue) and after (Red) exposure to AdipoRon. *B* and *D*: Summarized data from experiments similar to those in *A* & *C*, respectively. Data are shown as boxes and whiskers, with centerline as median value. Data are plotted in Box (75% of the data) and whiskers format (minimum to maximum), with centerline as median value. Data are compared to control responses with paired Student *t*-test. *, p < 0.05.

### ADPN receptor agonist AdipoRon dose-dependently inhibits BSM contraction force

As *Adpn^−/−^* mouse BSM exhibited profoundly diminished contraction force, we tested whether pharmacological modulation of ADPN signaling could regulate acute BSM contraction force. Interestingly, the selective ADPN receptor agonist, AdipoRon (33) dose-dependently inhibited EFS-induced BSM contractile force in both *wild-type* and *Adpn^−/−^* mice (Fig. 4*A*-*D*). AdipoRon inhibition of BSM contractile force was apparent at ∼1 µM and complete at ∼25 µM in both *wild-type* and *Adpn^−/−^*mice, indicating intact ADPN receptors in *Adpn^−/−^* mouse bladder. AdipoRon also dose-dependently inhibited BSM contractile force stimulated by carbachol-, by α,β-meATP and by KCl (Supplemental Fig. 2). These data reveal a novel function of AdipoRon in regulating acute BSM contraction force.

### AMPK signaling is crucial for BSM purinergic contraction force

AMPK is an energy sensor that regulates glucose and energy metabolism (34), and a well-known downstream effector of ADPN receptor activation. However, a role of AMPK signaling in ADPN receptor-mediated BSM contraction has not been reported. We examined BSM contractile responses to AMPK activators and inhibitors. As shown in Fig. 5*A* & *C*, the selective AMPK inhibitor Bay3827 partially inhibited EFS-induced BSM contraction. Interestingly, this inhibition seems more sensitive to lower-frequency than to high-frequency EFS-induced BSM contraction (Fig. 5*C*). As BSM purinergic contraction shares higher sensitivity to lower-frequency EFS, whereas high-frequency EFS-induced BSM contractility predominantly reflects muscarinic contraction, we suspected that AMPK inhibitor Bay3827 might specifically inhibit purinergic contraction of BSM. Indeed, in the presence of atropine to block muscarinic receptors while sparing purinergic signaling (Fig. 5*B*), Bay 3827 inhibited most of the purinergic contractility of BSM (Fig. 5*B* & *D*).

**Figure 5.**
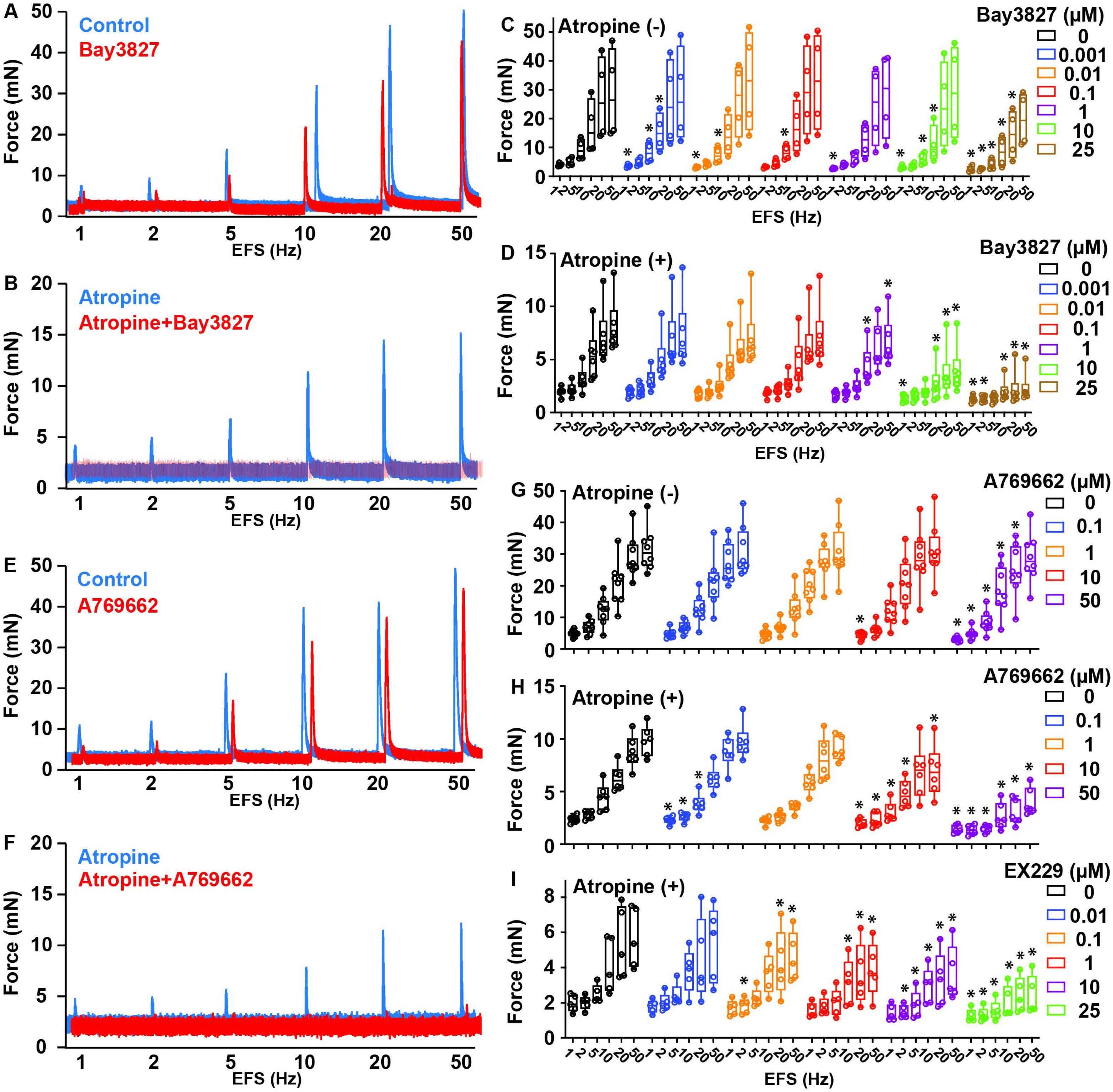
AMPK modulators inhibit mouse BSM purinergic contraction. *A*: Representative traces of *wild-type* BSM contraction in response to EFS (n=4) before (Blue) and after (Red) exposure to AMPK inhibitor Bay3827. *B*: Representative traces of atropine pre-treated *wild-type* male BSM contraction in response to EFS (n=6) before (Blue) and after (Red) exposure to AMPK inhibitor Bay3827. *C* and *D*: Summarized data from experiments in the formats of *A* & *B*. *E*: Representative traces of *wild-type* male BSM contraction in response to EFS (n=8) before (Blue) and after (Red) exposure to AMPK activator A769662. *F*: Representative traces of atropine pre-treated *wild-type* male BSM contraction in response to EFS (n=6) before (Blue) and after (Red) exposure to AMPK activator A769662. *F*: Representative traces of atropine pre-treated BSM contraction before (Blue) and after (Red) AMPK activator EX229 treatment, from male *wild-type* (n=5) mice in response to EFS. *G*, *H*, and *I*: Summarized data from experiments of the formats of *E* & *F*. Data are plotted in Box (75% of the data) and whiskers format (minimum to maximum), with centerline as median value. Student *t*-test. *, p < 0.05.

We further tested whether AMPK activation could impact acute BSM contraction force. To our surprise, AMPK activation also inhibited the purinergic contraction of BSM. We used a potent AMPK activator A769662 to test this possibility. As did AMPK inhibitor Bay3827, the potent AMPK activator A769662 partially inhibited EFS-induced BSM contractility, reflecting near-complete inhibition of the purinergic component of BSM contraction (Fig. 5*E*-*H*). The additional selective and potent AMPK activator EX229 similarly inhibited purinergic BSM contractility (Fig. 5*I*). These apparently paradoxical pharmacological data suggest that both activators and inhibitors of AMPK can inhibit purinergic contraction of BSM.

### *Adpn^−/−^* mice bladders exhibit altered expression of purinergic signaling molecules

To further understand the mechanism of impaired smooth muscle contractile function in *Adpn^−/−^* mice, we performed western blot studies to detect proteins of the signaling pathways regulating BSM contraction. Consistent with our functional data showing diminished EFS-induced BSM contractility (Fig. 3), we found expression levels of both CHRM3 receptor and P2X1 receptor to be significantly downregulated in *Adpn^−/−^* BSM (Fig. 6*A*, *B*, *G*, & *H*). The lack of purinergic contractility in *Adpn^−/−^* BSM prompted further interrogation of other purinergic signaling components in the bladder wall. Interestingly, BSM-expressed ATP-ADP-adenosine converting enzymes ENTPD1 and NT5E were both significantly upregulated in *Adpn^−/−^* bladders (Fig. 6*C*, *E*, *I*, & *K*). The interstitium-localized ENTPD2 and urothelium-expressed alkaline phosphatase (ALPL) were also significantly increased (Fig. 6*D*, *F*, *J*, & *L*) (35; 36). The profound changes of these purine nucleotide phosphatases could contribute to diminished purinergic contraction in *Adpn^−/−^*bladders.

**Figure 6.**
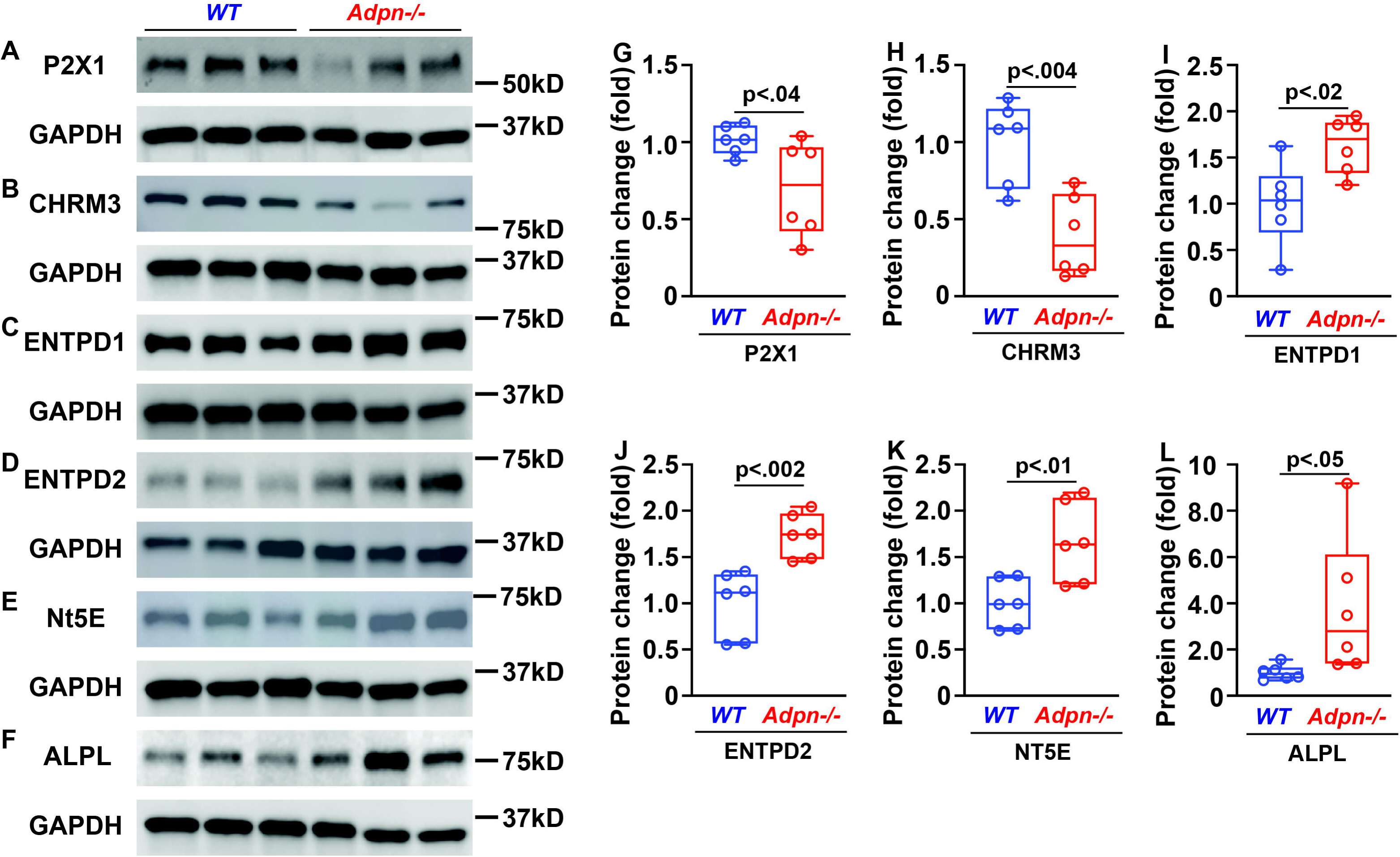
Alterations in purinergic signaling components correlate with diminished BSM purinergic contractile force in *Adpn^−/−^* bladder. (A-F) Western blots of P2X1, CHRM3, ENTPD1, ENTPD2, NT5E, and ALPL proteins from male *wild-type* and *Adpn^−/−^* mouse bladders (n=6). Quantitated data normalized to GAPDH (G-L) are plotted in Box (75% of the data) and whiskers format (minimum to maximum), with centerline as median value. Student *t*-test, *P* values above bars.

### *Adpn^−/−^* mice bladders exhibit altered proliferation and differentiation

ADPN signaling is known to regulate cell phenotype through transcription factors such as PPARα (26). Although the *Adpn^−/−^* mouse bladder weight did not differ from that of wild-type controls, the bladder/body weight ratio of *Adpn^−/−^*bladder was significantly lower than wild-type (Fig. 7*F*-*I*), suggesting altered cell proliferation and/or differentiation in *Adpn^−/−^* bladder wall. Indeed, numbers of Ki67-positive nuclei in the BSM layer of *Adpn^−/−^*mouse bladder were significantly lower than those found in wild-type controls (Fig. 7*A*, *B*, & *J*). Intriguingly, several smooth muscle marker proteins were significantly upregulated in *Adpn^−/−^* mice bladder, including α smooth muscle actin (αSMA), SM22, and smooth muscle myosin heavy chain (SMMHC) (Fig. 7*C*-*E*, & *K*-*M*), indicating an altered BSM differentiation and phenotype in *Adpn^−/−^* mice bladders potentially reflecting a futile compensatory response.

**Figure 7.**
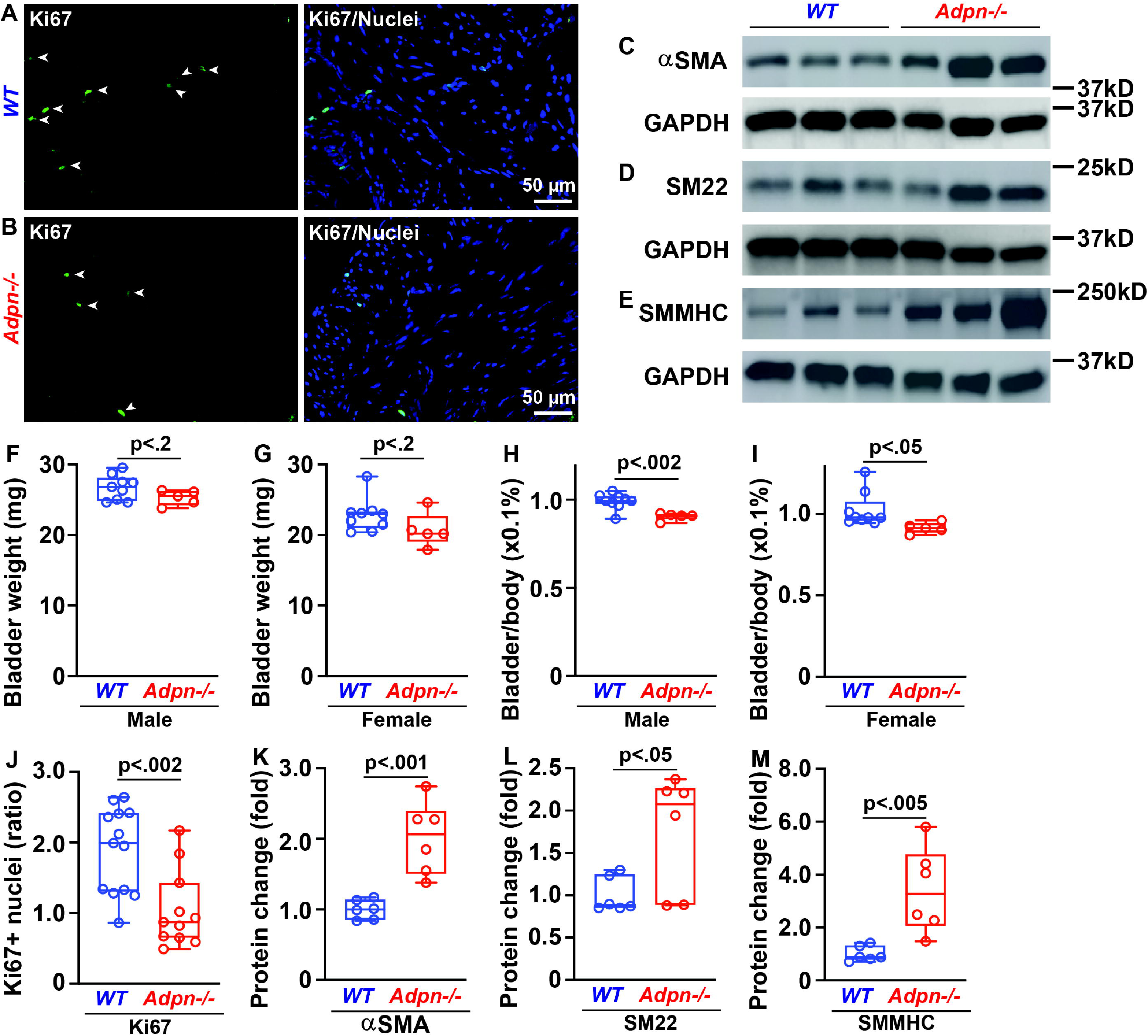
*Adpn^−/−^*BSM cells exhibit altered proliferation and differentiation phenotypes. A (*wild-type*) and B (*Adpn^−/−^*) bladder tissues immunostained for proliferation marker Ki67 (green, white arrowheads), colocalized with DAPI-stained nuclei (blue). *J*: Presents % of Ki67-positive cells in BSM layer (*wild-type* n=13; *Adnp^−/−^* n=11 sections). *C*-*E*: Western blots of smooth muscle markers αSMA, SM22, and SMMHC proteins in *wild-type* and *Adpn^−/−^* bladders (n=6). *K*-*M*: Summarizes GAPDH-normalized quantitated data. *F* - *I*: Bladder weights and bladder-to-body weight ratios of male (F, H) and female (G, I) *wild-type* (n=9) and *Adpn^−/−^* (n=5) mice. Data are plotted in Box (75% of the data) and whiskers format (minimum to maximum), with centerline as median value. Student *t*-test, *P* values above bars.

### *Adpn^−/−^* mice bladders exhibit altered downstream signaling

In addition to its importance in energy and metabolism homeostasis and insulin sensitization, ADPN signaling is also important in regulating gene transcription and protein/lipid synthesis, thereby impacting cell proliferation/differentiation (18; 37). Deletion of ADPN signaling in *Adpn^−/−^* bladder wall was confirmed by immunofluorescence and western blot (Supplemental Fig. 1). We have examined additional ADPN-relevant pathways in *Adpn^−/−^* bladders. As shown in Fig 8*A*, insulin receptor expression in *Adpn^−/−^* bladders did not differ from wild-type. AKT signaling is a major downstream pathway of insulin receptor activation important for glucose and lipid metabolism. Whereas total AKT in *Adpn^−/−^* bladders increased significantly, p-AKT (pSer473) was reduced compared to wild-type controls (Fig. 8*B*, *C*, *H*, & *I*). These data thus suggest impaired insulin signaling or potential insulin resistance in the bladder wall of *Adpn^−/−^* mice. Extracellular signal-regulated kinase (ERK) signaling is another pathway downstream of both ADPN receptors and insulin receptors, important for gene transcription and cell growth (38). *Adpn^−/−^*bladders exhibited decreased total ERK expression whereas p-ERK (pThr202 and pTyr204) was increased compared to wild-type (Fig. 8*D*, *E*, *J*, & *K*). AMPK expression in *Adpn^−/−^* bladder did not differ from that in wild-type (Fig. 8*F* & *L*).

**Figure 8.**
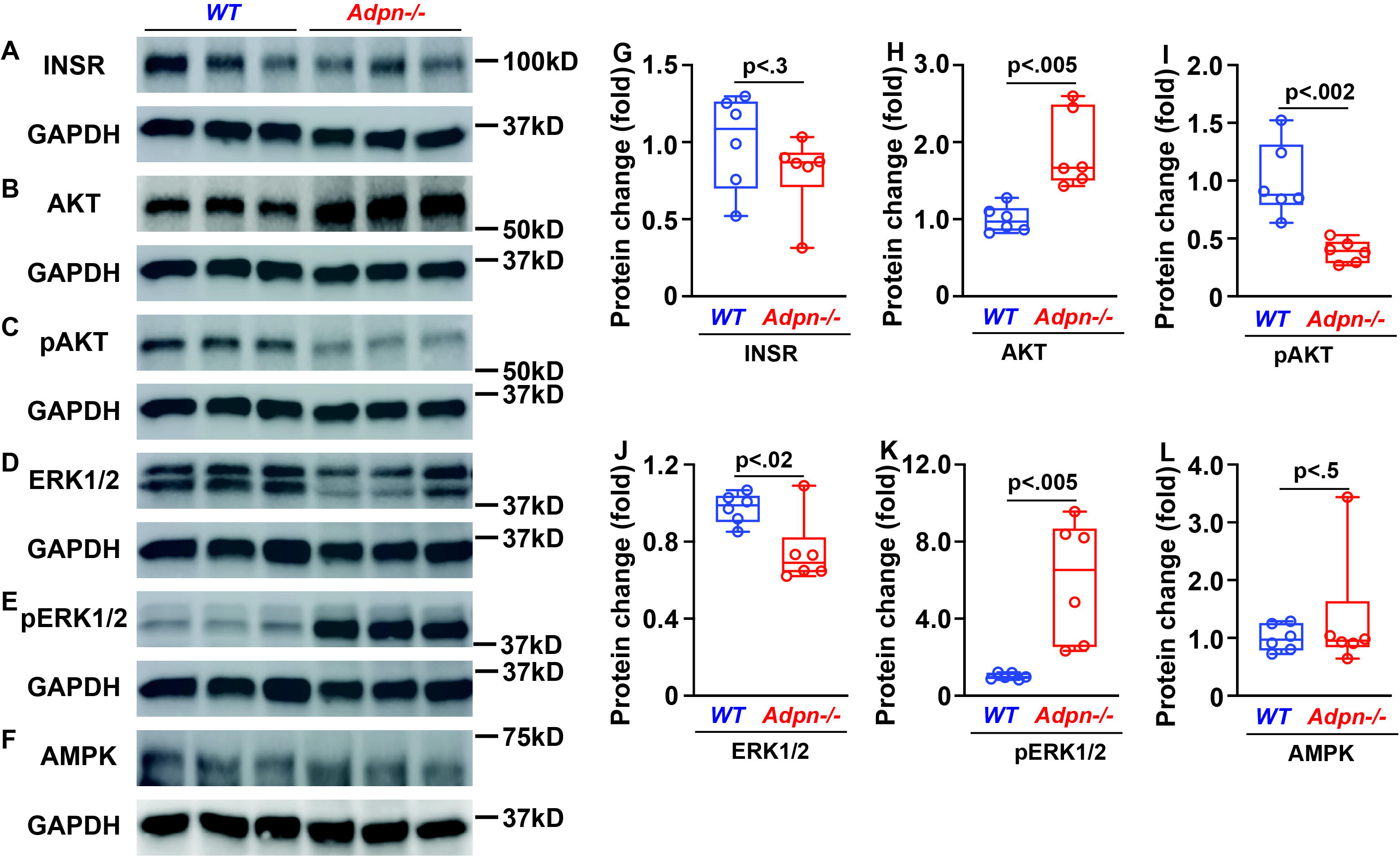
*Adpn^−/−^*BSM cells exhibit altered protein abundance in multiple downstream signaling pathways. *A*-*F*: Western blot of INSR, AKT, pAKT, ERK1/2, pERK1/2, and AMPK proteins from male *wild-type,* and *Adpn^−/−^*bladders (n=6). GAPDH was normalization control for quantitated data shown in *G*-*L*. Data are plotted in Box (75% of the data) and whiskers format (minimum to maximum), with centerline as median value. Student *t*-test, *P* values above bars.

## Discussion

The long-recognized clinical association of LUTS and obesity/metabolic syndrome has been little understood. Our study examined the hypothesis that adipokines are major contributors to the pathogenesis of LUTS associated with obesity/metabolic syndrome. We examined in particular the role of ADPN, the most abundant adipokine, in regulation of bladder function. Consistent with our hypothesis, *Adpn^−/−^* mice exhibited significantly increased voiding frequency, small voids, and decreased bladder compliance (Fig. 1 & 2). Interestingly, a recent clinical study reported that lower serum ADPN level was significantly associated with LUTS such as detrusor underactivity (DU). Lower serum ADPN levels were most strongly associated with decreased maximum flow rate (Qmax), decreased bladder voiding efficiency (BVE), and decreased bladder contractility index (BCI=Pdet @ Qmax + 5 Qmax, a quantitative value of bladder contraction and ability to sustain that contraction, where Pdet = detrusor pressure). Of note, ∼75% of patients with DU also exhibited detrusor overactivity (39). DU is defined as insufficient bladder contraction force and decreased voiding efficiency. Our mouse myography data corroborate this in the significantly diminished contractile force measured in *Adpn^−/−^* bladders (Fig. 3*A*-*C*). The bladder phenotype of *Adpn^−/−^* mice combined with the clinically reported inverse association of ADPN with LUTS together support an important role of ADPN in the regulation of bladder function. Levels of other adipokines including nerve growth factor (NGF), interleukins, and TNF-α have also been reported to correlate with OAB (40), suggesting contributions of additional molecular mechanisms to LUTS pathogenesis in association with obesity/metabolic syndrome.

BSM contraction is mediated by both muscarinic and purinergic signaling. We found that *Adpn^−/−^* bladder lacked a contractile response to purinergic activation (Fig.3*D*). Abundance of P2X1, a major purinergic receptor responsible for BSM purinergic contraction, was also significantly decreased in *Adpn^−/−^* bladder (Fig.6 *A*&*G*). In contrast, purine-converting and degrading enzymes in the *Adpn^−/−^* bladder wall, including ENTPD1/2, NT5E, and ALPL, were all upregulated (Fig.6 *C*-*F* & *I*-*L*). This upregulation is predicted to quickly eliminate purinergic agonists such as ATP and ADP from the *Adpn^−/−^*bladder wall. ATP and ADP stimulation of P2X1 and P2Y12 receptors activates contraction in human and rodent bladders (31). Thus, downregulated purinergic receptor activity combined with upregulation of purine nucleotide converting and degrading activities likely contribute to the diminished purinergic contraction force observed in *Adpn^−/−^* bladder.

Another novel finding of our study is the acute effect on BSM contraction by ADPN signaling. However, unlike the diminished contractile force in *Adpn^−/−^* bladder due to BSM phenotypic changes, ADPN receptor agonist AdipoRon acutely inhibited BSM contractile force in a dose-dependent manner within 10-15 minutes (Fig. 4 and Supplemental Fig. 2). This AdipoRon-mediated inhibition applied to tested contractile stimuli including EFS, carbachol, α,β-meATP, and KCl (Fig. 4 and Supplemental Fig. 2). The inhibitory effect was evident at 1 µM and near maximal at 10 µM, concentrations in agreement with the previously reported EC50 of AdipoRon (33). ADPN signaling exerts an acute vasodilatory effect by relaxing vascular smooth muscle cells (VSMC) through an undefined mechanism (41–43). ADPN can also hyperpolarize gastric smooth muscle resting membrane potential by increasing K^+^ current, thereby inhibiting excitation-contraction coupling (44). The mechanism by which ADPN inhibits acute smooth muscle contraction remains controversial. However, our observation of acute inhibition of smooth muscle contraction by AdipoRon appears unprecedented in the literature as of the time of writing. ADPN signaling has been reported to activate numerous intracellular signaling pathways, including the phospholipase C inositol 1,4,5-trisphosphate pathway regulating sarcoplasmic reticulum calcium release in skeletal muscle. A similar pathway mediates muscarinic contractility in BSM cells, but cannot be easily reconciled with our observation that AdipoRon inhibits BSM contraction. A more thorough mechanistic understanding of ADPN action in BSM may require studies of mouse models of selective ADPN receptor subtype knockout.

Interestingly, pharmacological inhibition of AMPK signaling specifically blocked acute purinergic contraction of BSM (Fig.5 *A*-*D*), consistent with the diminished purinergic contractility observed in *Adpn^−/−^* BSM (Fig. 3*D*). Paradoxically, pharmacological activation of AMPK signaling also acutely blocked purinergic contraction of BSM (Fig.5 *E*-*I*). This latter unexpected data was consistent with reported ADPN effects on vasodilation and gastric smooth muscle relaxation (41; 44), and with our AdipoRon data (Fig. 4 and Supplemental Fig. 2). AMPK is an energy-sensing master kinase known to modulate the activity of >100 proteins. AMPK activation by low-potency agonist AICAR (5-aminoimidazole-carboxamide ribonucleoside) partially inhibited aortic smooth muscle contraction (45). AMPK signaling has also been shown to inhibit tonic vascular smooth muscle contraction by direct AMPK phosphorylation of myosin light chain kinase (MLCK) at Ser85, preventing MLCK activation by blocking its calmodulin binding site (46). Perhaps AMPK regulates a finely balanced energy state required to achieve maximal contractile force in response to acute stimulation by neurotransmitter and autocrine agonists. This balanced resting state may reflect the well-known optimized pre-stretched smooth muscle length required to generate subsequently stimulated maximal contractile force, such that either over-stretched or relaxed basal states will generate reduced contractile force (47).

ADPN signaling has been shown to regulate both proliferation and differentiation of VSMC, each important for the maintenance of normal VSMC contractile phenotype (48–50). The reduced cell proliferation in *Adpn^−/−^* bladder (Fig. 7) might reflect decreased p-AKT signaling (Fig. 8*C*). In contrast, smooth muscle biomarkers αSMA, SM22, and SMMHC (Fig 7) were all increased in abundance, with correspondingly increased p-ERK (Fig. 8*E*). We speculate that reduced proliferation in *Adpn^−/−^* bladder (Fig. 7*A*, *B*, *F*-*J*) may reflect a hypertrophic response mediated through p-ERK signaling to compensate deficient BSM contractile function (38). The data together indicate considerable complexity of ADPN signaling in BSM.

Our study has several important limitations. Although circulating ADPN derives mainly from adipocytes, BSM also expresses ADPN locally (16). The relative importance or differential functions of adipocyte-derived and BSM-expressed ADPN in regulating bladder function has not been addressed. The specific ADPN receptors involved and details of receptor-specific downstream signaling pathways are yet to be studied. Smooth muscle-specific ADPN receptor knockout animal models are currently under development in our laboratory to further address these central questions.

In summary, we have discovered a bladder phenotype in *Adpn^−/−^*mice that recapitulates symptoms and urodynamics observed in humans with LUTS associated with obesity/metabolic syndrome. We have also discovered that ADPN signaling plays a profound role in regulating BSM contractility and urodynamics. Disruption of ADPN signaling leads to both metabolic and purinergic signaling pathway changes in the bladder, which we propose may play important roles in the pathogenesis of LUTS associated with obesity/metabolic syndrome.

## Author Contributions

W.Y. conceived and supervised the project, performed cystometrograms and myography, analyzed data and wrote the manuscript. Z.L. performed histological and immunostaining & imaging, Western blot, and analyzed data. A.W. performed voiding spot assays. S.R. and S.L.A. contributed to the conception and interpretation of data, and revised the manuscript. All coauthors critically reviewed the manuscript, discussed ideas and results, and contributed to the manuscript. W.Y. is the guarantor of this work and, as such, has full access to all the data in the study and takes responsibility for the integrity of the data and the accuracy of the data analysis.

