## Supplemental Information for "Adiponectin Signaling Regulates Urinary Bladder Function by Blunting Smooth Muscle Purinergic Contractility"

Supplemental Figure 1. Adiponectin protein is absent from *Adpn^-/-^* mice BSM cells. *A*: *Wild-type*) and *B*: *Adpn^-/-^* bladder tissues immunostained with anti-adiponectin antibody (green), localized in BSM layer (*A*) and completely absent from *Adpn^-/-^* bladder tissue (*B*). Nuclei were labeled with DAPI (blue). *C*: Western blot of adiponectin protein detected a specific band above 25KD in *wild-type* bladder absent from *Adpn^-/-^* bladder (n=6), with GAPDH loading control.

Supplemental Figure 2. Adiponectin receptor agonist AdipoRon inhibited mouse BSM contractile force in a dose-dependent manner. Representative *wild-type* male BSM contraction traces in response to carbachol (*A*), α,β-meATP (*C*), and KCl (*E*), before and after exposure to AdipoRon. *B*, *D*, and *F*: Summarized data corresponding to experiments in formats of panels *A*, *C* and *E* (n=4-8). Data show individual symbols and line plots for each sample. Paired student *t*-test, *P* values above data points. *, P<0.05.

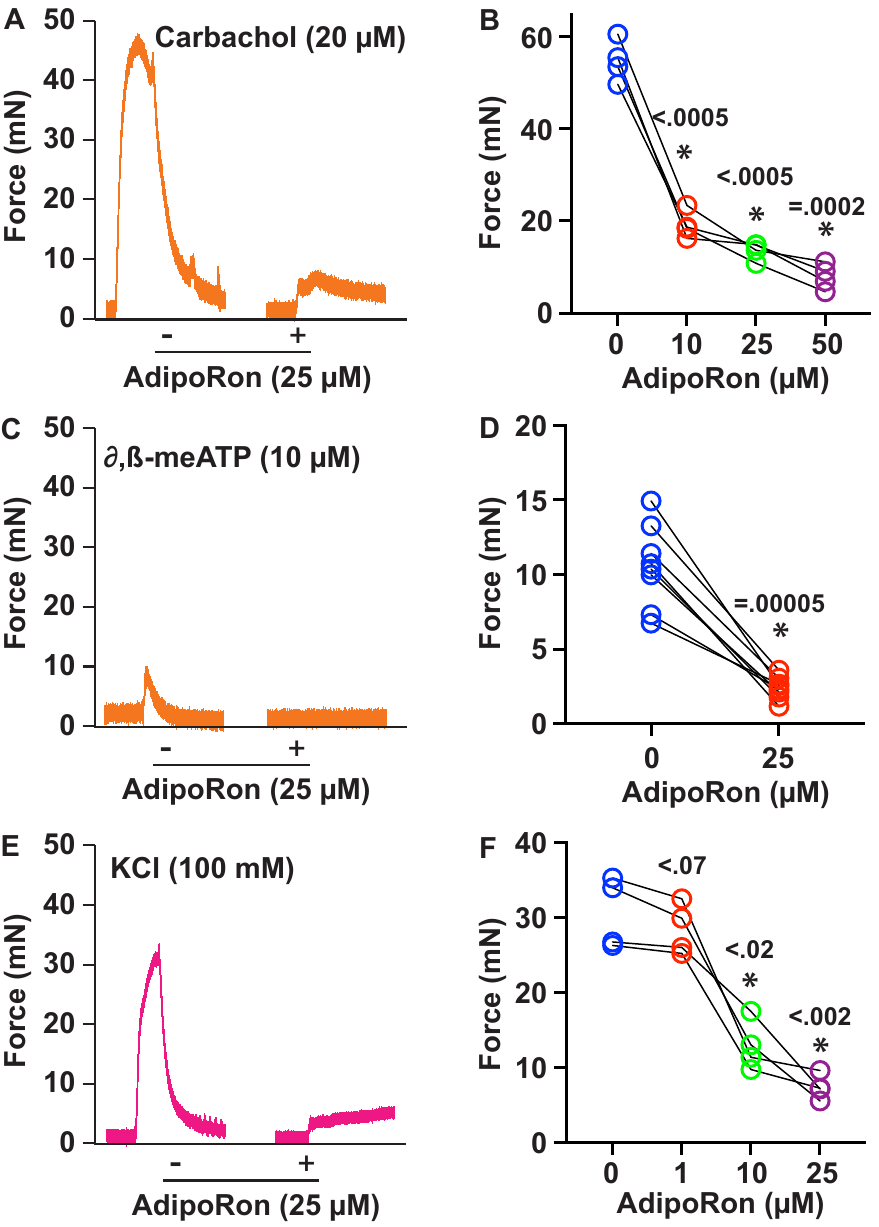

Supplemental Table 1. Antibody information

| Antibody | Company | Catalog | Host | Application |
| --- | --- | --- | --- | --- |
| Adiponectin | R&D system | AF1119 | Goat | WB, IF |
| Ki67 | Invitrogen | 14-5698-82 | Rat | IF |
| Alkaline Phosphatase | R&D system | AF2910 | Goat | WB |
| ENTPD1 | Invitrogen | MA5-32707 | Rabbit | WB |
| ENTPD2 | R&D system | AF5797 | Sheep | WB |
| CHRM3 | Invitrogen | PA585322 | Rabbit | WB |
| P2X1 | Alomone laboratory | APR-001 | Rabbit | WB |
| αSMA | Cell Signaling Technology | 19245S | Rabbit | WB |
| TAGLN | Cell Signaling Technology | 40471S | Rabbit | WB |
| SMMHC | Proteintech | 21404-1-AP | Rabbit | WB |
| 5'-Nucleotidase | R&D system | MAB44881 | Rat | WB |
| Insulin Receptor β | Cell Signaling Technology | 23413S | Rabbit | WB |
| p-AKT | Cell Signaling Technology | 9271T | Rabbit | WB |
| AKT | Cell Signaling Technology | 9272S | Rabbit | WB |
| p-Erk1/2 | Cell Signaling Technology | 9101S | Rabbit | WB |
| Erk1/2 | ABclonal | A4782 | Rabbit | WB |
| AMPK-α | Cell Signaling Technology | 2532S | Rabbit | WB |
| GAPDH | ABclonal | A19056 | Rabbit | WB |
